# Oral microbial signatures of head and neck cancer patients with diverse longitudinal oral mucositis severity patterns

**DOI:** 10.1101/2025.07.15.665024

**Authors:** Saritha Kodikara, Jiadong Mao, Erin Marie D. San Valentin, Kim-Anh Do, Cielito C. Reyes-Gibby, Kim-Anh Lê Cao

## Abstract

**Background:** Oral mucositis is a painful complication commonly observed in head and neck cancer patients receiving cancer treatment. Emerging evidence suggests that changes in the oral microbiome can contribute to oral mucositis development, making microbial signatures potential targets for therapeutic inter-ventions. This study aimed to: (1) characterize longitudinal microbial patterns of oral mucositis severity among head and neck cancer patients; (2) determine clinically relevant patient clusters based on oral mucositis severity trajectories; and (3) identify microbial signatures specific to these clusters.

**Results:** We derived a calibrated oral mucositis score by applying non-negative sparse principal component analysis to seven oral mucositis related symptom ratings, using longitudinal microbiome data from 140 head and neck cancer patients. Functional data analysis and hierarchical clustering identified three distinct patient clusters with differing microbial trajectories of oral mucositis progression. One cluster exhibited patients with a rapid increase in oral mucositis severity following treatment initiation, while the other clusters displayed more gradual increase. Demographic comparisons revealed significant differences in age and weight distributions between clusters, with older, lighter patients more common in clusters experiencing more gradual oral mucositis progression. Partial least squares knockoff analysis identified cluster-specific microbial signatures: notably, *Prevotella spp.* positively associated with calibrated oral mucositis score across all clusters, while *Alloprevotella* (*Alloprevotella0302*) was significantly enriched only in patients experiencing rapid oral mucositis progression. Conversely, genera associated with oral health, including *Haemophilus, Rothia*, and *Actinomyces*, were negatively correlated with calibrated oral mucositis score.

**Conclusions:** Distinct trajectories of oral mucositis scores in head and neck cancer patients are linked to specific oral microbial profiles and demographic factors. The identification of cluster-specific microbial profiles highlights the potential for microbiome-targeted interventions to manage oral mucositis severity. While most taxa were cluster-specific, *Prevotella* consistently ranked among the top taxa positively associated with the calibirated oral mucositis score across clusters, suggesting it may not differentiate between patient groups but rather reflects overall disease severity.

## 1 Introduction

Head and neck squamous cell carcinoma (HNSCC) affects the upper aero-digestive tracts and often leads to symptoms such as eating difficulty, swallowing pain and ear pain (Johnson et al, 2020). Oral mucositis (OM) commonly develops in HNSCC patients, often as a side effect of radiotherapy and chemotherapy (Pulito et al, 2020). OM is characterized by inflammation and ulceration in the mouth, which exacerbates the symptoms experienced by HNSCC patients (Bell and Kasi, 2025). Although various strategies targeting the patients’ oral microbiome have been explored, an approved and effective OM treatment is still lacking (Zhang et al, 2024).

OM development involves a complex interplay of immune dysregulation, tissue damage, and shifts in the oral microbiome. While the microbiome normally helps maintain oral health, it becomes disrupted during cancer treatment, contributing to OM progression and severity (San Valentin et al, 2023; Li et al, 2020; Zhang et al, 2024; Reyes-Gibby et al, 2020). Despite numerous intervention strategies that have been explored, an effective and widely approved treatment for OM remains elusive. Thus, understanding the microbial dynamics associated with OM development and progression may provide valuable insights into potential therapeutic targets and preventive measures.

However, tracking OM development and the microbial dynamics is challenging due to the lack of a robust OM severity score, summarizing main symptoms of OM, and the sparse and noisy nature of longitudinal microbiome data. Based on a thorough analysis of data in Zhang et al (2024), we move well beyond their original descriptive work by (i) deriving a statistically calibrated composite oral mucositis score that fuses pain, erythema, ulceration, and functional impairment into a single clinically meaningful continuum; (ii) applying state-of-the-art functional data analysis to those trajectories to define data driven patient groups with shared symptom dynamics; and (iii) deploying the PLS knockoff procedure to pinpoint microbial communities that robustly distinguish the resulting trajectory groups (Yang et al, 2025). This integrated analysis delivers mechanistic insight that could not be achieved through conventional analyses and provides a transferable template for linking longitudinal symptom burden with the microbiome in oncology cohorts.

## 2 Materials and methods

### 2.1 Data

Zhang et al (2024) enrolled 145 HNSCC patients and generated a rich longitudinal data set – 28 self-reported symptoms rated on a 0–10 scale at up to 14 time points per patient (median = 8; Fig. 1) using the MDASI-H&N instrument (Rosenthal et al, 2007), together with 16S rRNA V4 profiles comprising 982 OTUs.

**Fig. 1.**
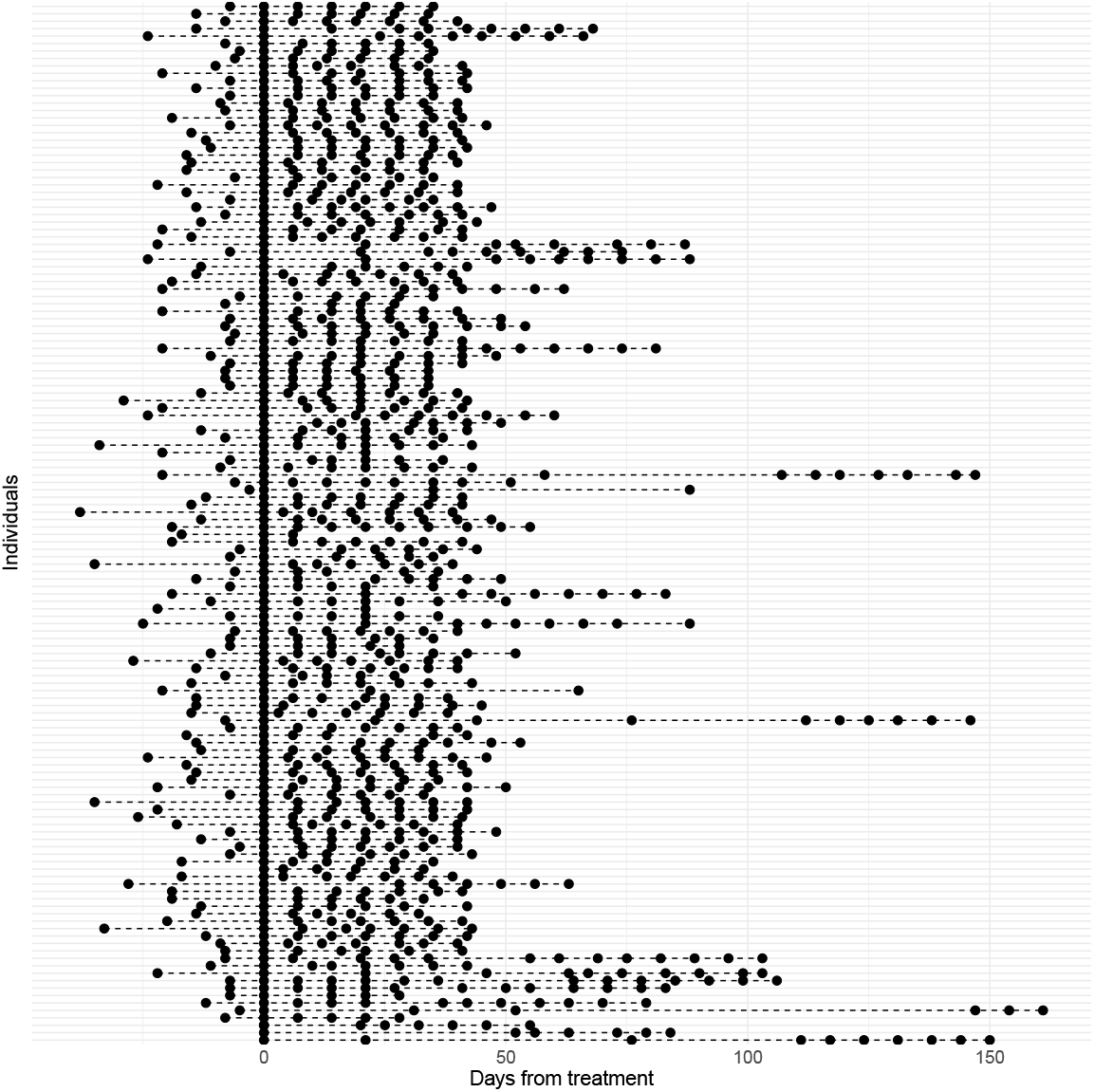
The swimmer plot of 140 patients highlights marked variability in observation timing and total study duration, posing challenges for standard analyses. This irregularity necessitates flexible modelling approaches, such as functional data analysis and linear mixed models, which can accommodate unbalanced and irregularly spaced longitudinal data.

We refined this resource in three key ways before any downstream modeling. (1) Cohort anchoring: we retained only the 140 patients with complete baseline records, ensuring every trajectory begins from a common reference point. (2) Stringent feature curation: OTUs whose cumulative counts fell below the 5% percentile were discarded, paring the feature space from 982 to 147 high-confidence taxa and removing noise from sequencing artifacts and vanishingly rare organisms. (3) Compositional validity: the filtered count matrix was centered-log-ratio transformed, converting relative abundances to Euclidean coordinates suitable for multivariate analysis. These pre-processing steps enabled to conduct the subsequent functional clustering and knock-off inference.

### 2.2 Methods

#### Data related challenges

Longitudinal studies introduce several methodological hurdles. First, there are still no widely adopted tools for clustering patients by irregular symptom trajectories or for detecting OTUs whose relationships with oral-mucositis (OM) scores differ between patient groups (Kodikara et al, 2022; Peterson et al, 2024). In our cohort, each patient’s symptom profile forms a multivariate time series recorded at uneven intervals within individual study windows (Fig. 1); clustering methods tailored to such unbalanced data remain scarce (Aghabozorgi et al, 2015; Ghaderi et al, 2024). Second, standard differential-abundance analyses test only for shifts in mean OTU abundance between groups (Peterson et al, 2024). They cannot reveal taxa whose association with a continuous outcome such as the OM score varies across clusters, thereby missing group-specific microbial drivers of disease severity.

#### Calibrated OM score summarizing all relevant symptoms

We derived the calibrated OM score based on patients’ self-reported symptom ratings collected at multiple time points. To facilitate this, we first collapsed the time variable to obtain a matrix, where each row corresponds to a patient at a certain time point, and each column is a symptom. The values in the matrix are ratings with integer values ranging from 0 (mild) to 10 (severe), representing severity of a given symptom at a given time point.

To summarize patients’ symptom ratings, we conducted non-negative sparse principal component analysis (NSPCA, Sigg and Buhmann 2008) of the double-centered data (data was centered by rows first, and then by columns). The double centering ensured that symptom ratings from different patients at different time points were directly comparable (Marron and Dryden, 2021). Compared to conventional PCA, NSPCA yields non-negative and sparse principal component (PC) loadings, often improving biological interpretability.

We selected PC1 score as the calibrated OM score since PC1 showed a strong positive correlation with the OM score previously defined by Zhang et al (2024) (Fig 2A) and also captured a large amount of variation across seven symptoms related to OM (Fig 2B), thus offering a comprehensive summary of oral mucositis severity.

**Fig. 2.**
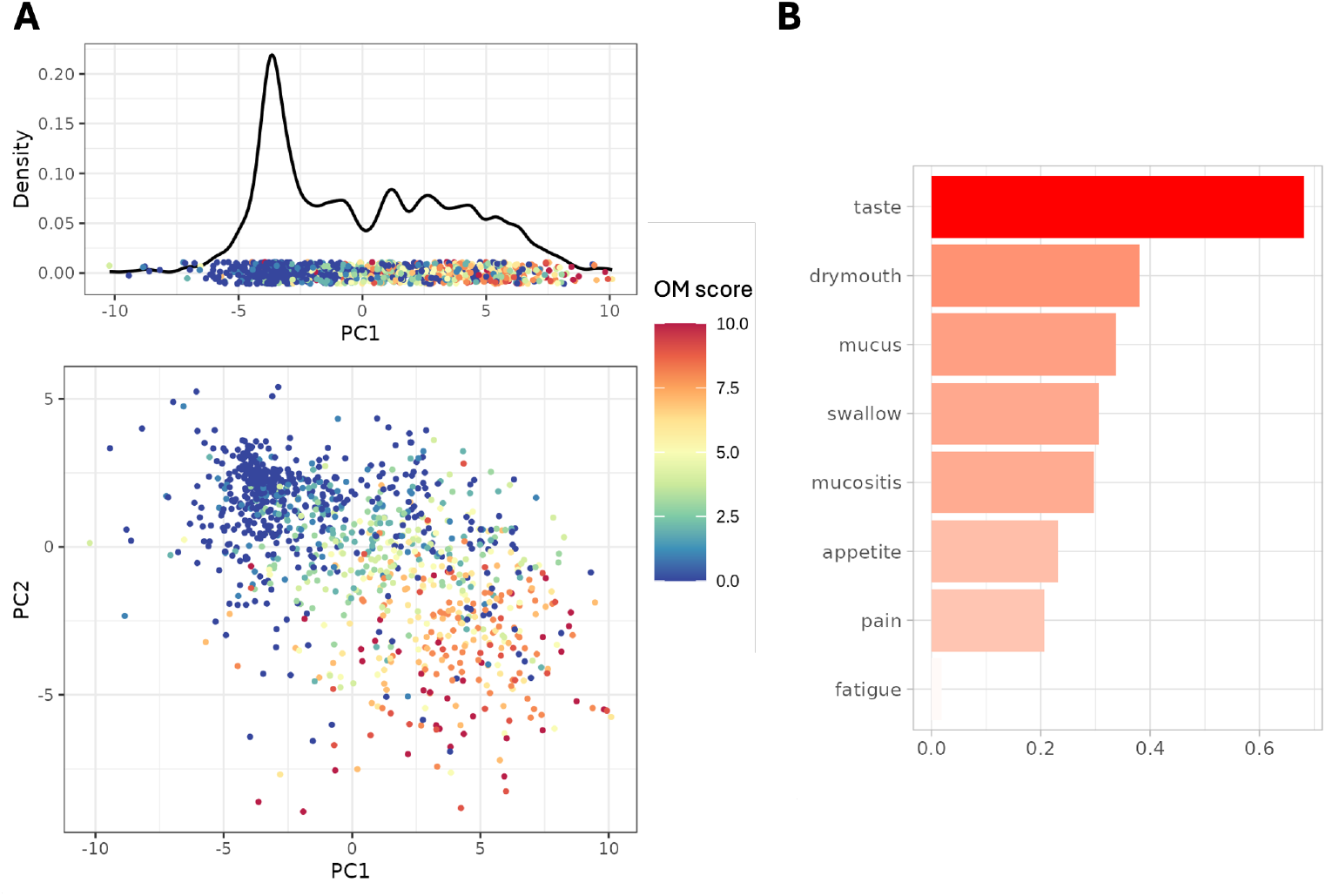
**A:** First and second principal components (PCs) from nonnegative sparse PCA based on all patients’ longitudinal ratings of their symptoms. PC1 is positively correlated with the calibrated OM score, which was used by Zhang et al (2024) as calibrated OM score, while PC2 is negatively correlated with calibrated OM score. **B:** Variable loadings from PC1 confirming PC1 is correlated with a comprehensive range of OM symptoms.

#### Identifying patient clusters according to calibrated OM score to reveal shared disease progression patterns

After obtaining a calibrated OM score for each patient at their respective observational time points, we used a functional data analysis approach to cluster the patients according to the longitudinal patterns of their calibrated OM scores (Marron and Dryden, 2021).

For the *i*th patient, we linearly interpolated their observed calibrated OM scores at discrete time points (Fig 1). This gave a continuous OM score function *f*_*i*_(*t*) for *t* ∈ [*τ*_*i*_, *T*_*i*_], where *τ*_*i*_ and *T*_*i*_ represent the *i*th patient’s baseline and discharge time points, respectively (Fig 1). To preserve each patient’s original observation window, we limited *f*_*i*_ to their actual time range.

We then calculated the pairwise distances between the patient’s calibrated OM score functions as follows. For patients *i* and *j*, we define their distance *d*_*i,j*_ as

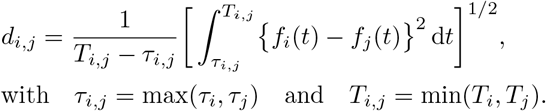

That is, *d*_*i,j*_ is defined as the mean squared distance between *f*_*i*_ and *f*_*j*_ over the their shared time interval [*τ*_*i,j*_, *T*_*i,j*_].

With the pairwise distances defined, we applied hierarchical clustering to the distance matrix to identify clusters of patients with comparable disease dynamics. The number of clusters was chosen to be 3 based on the gap statistic (Tibshirani et al, 2001).

#### Identifying underlying patient characteristics that are driving patient clusters

To investigate whether specific clinical features differed significantly between patient clusters, we first visualized marginal density plots for relevant variables such as age, weight, and BMI. In addition to visual differences between clusters, we also performed two-sample t-tests. Assumptions of normality and homogeneity of variance were checked prior to conducting these tests.

#### Identifying microbial features associated with calibrated OM score

To identify microbial features associated with calibrated OM scores, we used a novel variable selection method, partial least squares knockoff (PLSKO, Yang et al 2025). PLSKO is a knockoff variable selection method based on the knockoff framework proposed by Barber and Candes (2015) and PLS regression (Wold, 1966; Lê Cao and Welham, 2021). We chose a knockoff-based approach because many commonly used feature selection methods such as lasso (Tibshirani, 1996) and sparse PLS (Lê Cao et al, 2011) do not offer rigorous control over false discoveries, particularly in highdimensional and highly correlated settings, typical of microbiome data. In contrast, knockoff methods provide a statistically principled framework that guarantees false discovery rate control under minimal assumptions, making them especially well-suited for identifying robust microbial signatures (Peterson et al, 2024). We first applied PLSKO to the whole dataset to identify OTUs correlated with the calibrated OM score across all patients. We subsequently applied PLSKO to each individual cluster of patients to uncover cluster-specific OTUs.

#### Identifying patient characteristics that are associated with microbial abundance

To investigate associations between microbial abundance and patient characteristics (such as age, height, and weight) as well as study-related factors (such as time point and patient cluster membership), and to account for repeated measurements and inter-individual variability, we applied linear mixed models (LMMs). LMMs extend standard linear models by including both fixed effects and random effects, making them particularly suitable for repeated measurements. In our models, age, height, weight, time point, and cluster were treated as fixed effects, while patients were included as random effects to account for dependencies in the data. We fitted a separate LMM to each OM score-associated taxon identified by PLSKO, using CLR-transformed microbial abundance as the response variable.

The resulting regression coefficients from these models represent the estimated effect size of each fixed effect on microbial abundance. The sign and magnitude of each coefficient *β* represent the direction and strength of the association for a given covariate (e.g. time point) while keeping other covariates constant. We then plotted *sign*(*β*) *×*− log(*β*) in a heatmap to visualize the most significant drivers of microbial abundance.

## 3 Results

### Interpretable calibrated OM score reveals different longitudinal patterns of OM symptom progression

The longitudinal calibrated OM scores for individual patients showed a nonlinear upward trend (Fig 3A). In particular, the curve remained relatively stable before the start of treatment (Day 0), followed by a sharp increase until Day 25 before reaching a plateau. The overall trend shown in Fig 3A suggests that cancer treatment led to a substantial increase in OM symptom severity. While all patients exhibited increasing trend, we identified three distinct trajectory clusters (see Section 2). For example, the calibrated OM scores for patients in cluster 2 increased more sharply than those in clusters 1 and 3 (Fig. 3B).

**Fig. 3.**
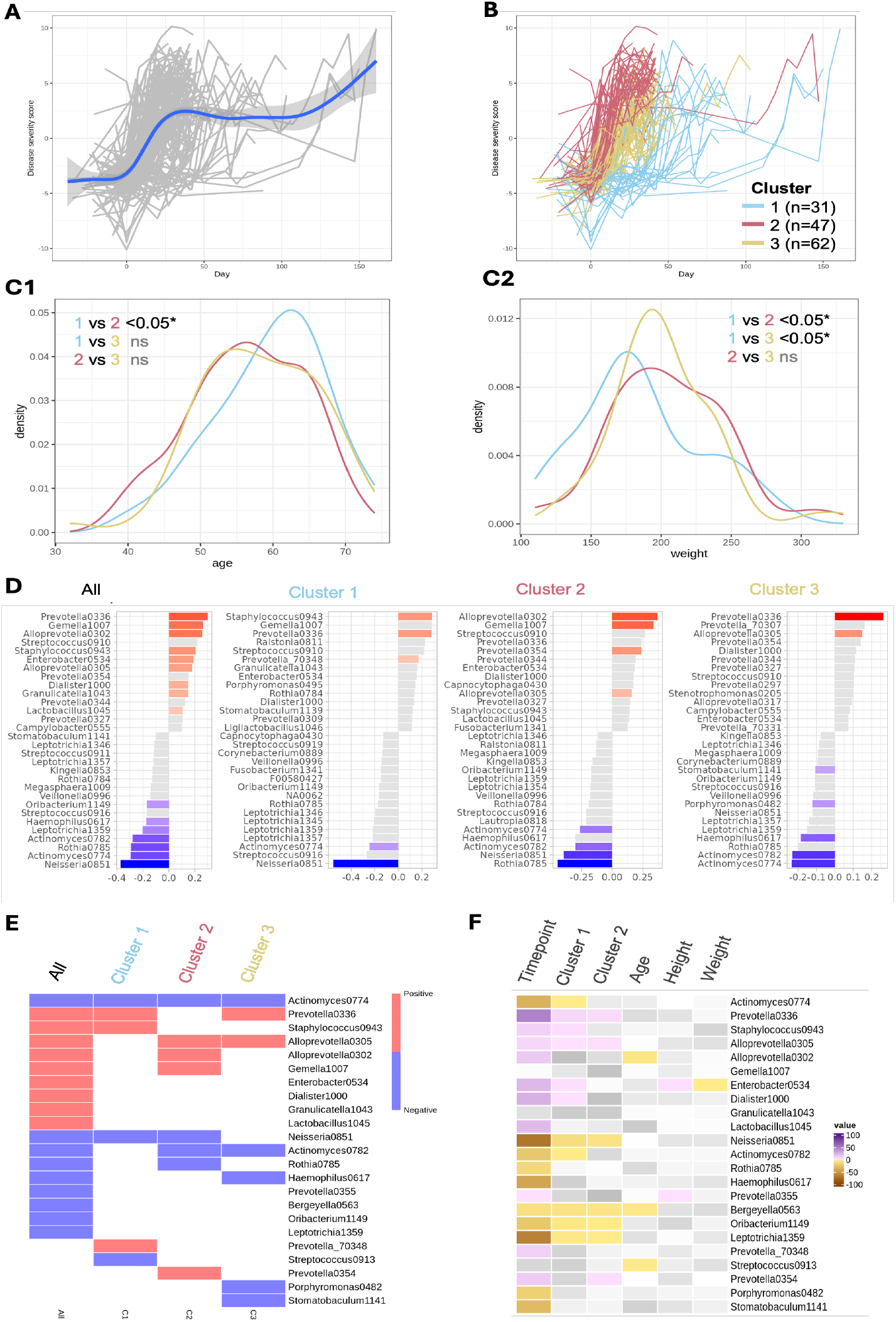
**A:** Calibrated OM score as a function of time (days since start of treatment). Each curve represents a patient, with mean curve indicated in blue and confidence band for the mean curve in gray. **B:** We clustered patients according to their calibrated OM score curves, resulting in three clusters. **C:** Probability density plots illustrating differences in age (**C1**) and weight distributions across patient clusters (**C2**), p-values from two-sample t-tests. We observe a significant difference in both age and weight distributions between clusters 1 and 2. **D:** Loading plots showing OTUs most strongly associated with calibrated OM scores among different patient clusters. Colored bars represent OTUs selected by PLSKO, i.e. OTUs with significant association with calibrated OM scores. **E:** Patient cluster-specific microbial features based on D, colored according to the sign of their correlation with calibrated OM scores within each cluster. **F:** Significant regression coefficients in the linear mixed models (LMM). Each row corresponds to a LMM with one of the significant taxa in **(E)** as response and time point, patient cluster membership and selected demographic features as predictors. Significant predictors are colored according to their coefficient values; non-significant ones are in gray.

### Patient cluster-specific demographic characteristics

We compared demographic features of patients in each pair of patient clusters (see Table 1). Patients in cluster 1 differed significantly in age and weight compared to those in cluster 2 (Fig. 3C1&C2). Cluster 1 patients, who exhibited a slower and more gradual increase in calibrated OM score (Fig 3B), were also older and had lower body weight than those in other clusters.

**Table 1.**
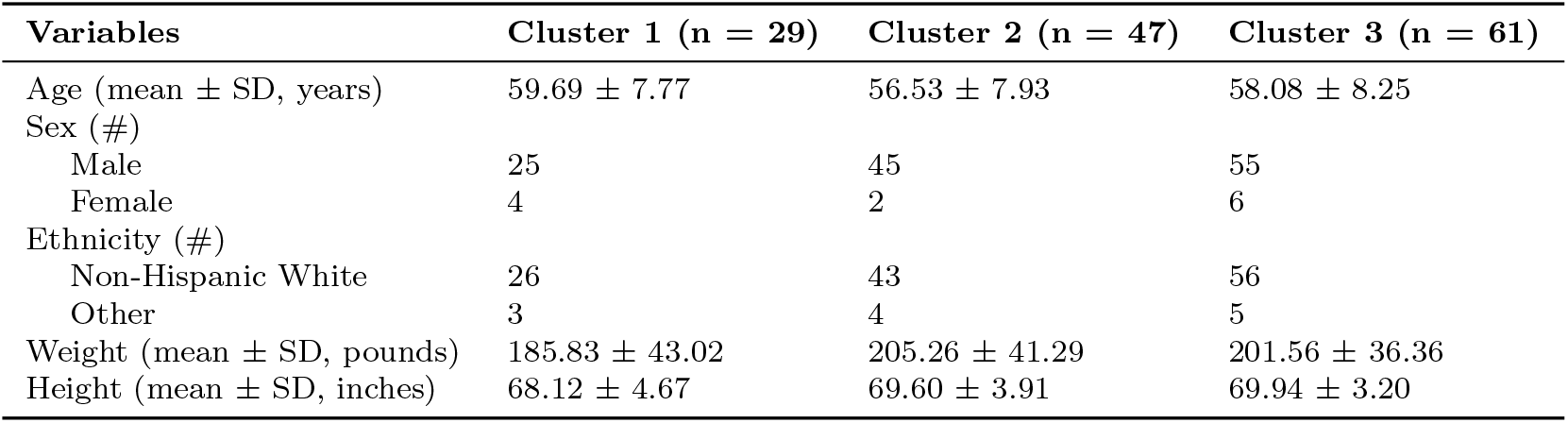
Demographic characteristics of patient clusters.

### Patient cluster-specific microbial signatures associated with calibrated OM score

The PLS regression analysis identified microbial OTUs that were most strongly associated with the calibrated OM score in different patient clusters (Fig 3D-E). The genus *Prevotella* consistently ranked among the top five OTUs positively associated with calibrated OM score across all clusters. This indicates an overall positive association with the calibrated OM score. The role of *Prevotella* was also highlighted in Zhang et al (2024). However, the original study did not explore OTUs negatively associated with the calibrated OM score. Through our analysis, we found that many of the bacterial genera negatively associated with our calibrated OM score, e.g. *Haemophilus, Rothia, Actinomyces*, has been found abundant in the healthy oral microbiome (Bik et al, 2010).

Using PLSKO for variable selection, we identified cluster-specific microbial signatures. For instance, an OTU belonging to genus *Alloprevotella* (*Alloprevotella0302*) was only found significantly associated with calibrated OM score in cluster 2 (Fig 3D). Cluster 2 exhibited the sharpest increase in the calibrated OM score. Importantly, *Alloprevotella* has also been detected in patients with oral dysbiosis (Devaraja and Aggarwal, 2025). Additionally, cluster 1 showed the largest difference in negative loadings (Fig 3D), further highlighting its distinct behavior between the three clusters.

Of note, we could not link OTUs associated with lower calibrated OM score with bacterial genera found in existing probiotic or prebiotic products manufactured by, e.g., ProBiora, Gallinée, Riven and MouthFlora. These products typically contain bacteria from genera such as *Streptococcus, Lactobacillus* and *Bifidobacterium* which are found abundant in healthy *gut* microbiome but not in healthy *oral* microbiome. This observation underscores the need to develop oral prebiotic and probiotic formulations specifically tailored to the oral microbiome, enabling therapeutic modulation of the microbiome in patients with HNSCC.

### Patient characteristics linked to microbial signatures associated with OM severity

Fitted LMMs for the significant taxa identified in Fig 3D-E revealed key drivers of microbial variation. In particular, time and patient cluster membership significantly explained the variation in most of the selected taxa. In particular, taxa positively correlated with calibrated OM score tended to have increased abundance as treatment progressed, whereas taxa negatively correlated with calibrated OM score tended to decrease over time. In addition, taxa that increased (or decreased) over time were typically more (or less) abundant in patient clusters 1 or 2, highlighting a clear link between temporal dynamics and cluster-specific microbial profiles. In contrast, patients’ age, height and weight were mostly not significant in explaining the variation of selected taxa.

## 4 Discussion

This study provides new insights into the longitudinal progression of OM in patients with HNSCC by integrating symptom trajectories and microbiome profiles. By clustering patients based on their OM score trajectories and identifying group-specific microbial signatures, we highlight potential microbial contributors to disease severity and variability in patient experience.

Our findings suggest that the three patient clusters reflect clinically meaningful differences, and that microbial dysbiosis may be associated with the progression of OM in patients with HNSCC. The distinct age and weight profiles observed among clusters 1, 2, and 3 suggest that these demographic factors may influence disease trajectory patterns. While younger patients tend to develop OM due to the more rapid rate of basal cell turnover, lesions also heal more quickly in younger patients (Bell and Kasi, 2025; Sonis, 1998). Older age becomes a risk factor for severe OM possibly due to insufficient DNA repair (San Valentin et al, 2023; Balducci and Extermann, 2000; Lionel et al, 2006). These findings are consistent with prior observations by Li et al (2020), who reported that a body weight loss exceeding 5% during chemoradiation in nasopharyngeal carcinoma (NPC) patients predicted severe OM. Interestingly, another study identified pre-treatment overweight status as an independent risk factor for OM severity in NPC patients (Li et al, 2020).

Microbial shifts also varied across the patient clusters. Severe OM cases exhibited a higher prevalence of inflammation-associated taxa such as *Staphylococcus, Fusobacterium*, and *Prevotella*. These organisms are consistently implicated in the onset of severe OM in HNSCC patients (Zhang et al, 2024; Reyes-Gibby et al, 2020). Particularly, *Prevotella* and *Fusobacterium* imbalances have been previously linked to oral dysfunctions such as halitosis (Carda-Diéguez et al, 2022; Hampelska et al, 2020) and periodontitis (Lima et al, 2023), with its Gram-negative properties linked to periodontal tissue destruction. Commensal bacteria such as *Neisseria, Rothia, Oribacterium, Haemophilus, Leptotrichia*, and *Actinomyces* were also found to be significantly depleted in patients with severe OM, suggesting a loss of the protective microbial communities coupled pathogenic overgrowth. *Rothia* and *Neisseria* are consistently found at higher levels in healthy individuals (Könönen et al, 2022; Pan et al, 2023).

Cluster-specific analyses also revealed notable differences in microbial composition. Cluster 1 showed a significant enrichment of *Staphylococcus* and *Prevotella*, and a concurrent depletion of *Neisseria* and *Leptotrichia*. This specific pattern may contribute to the observed longer and gradual increase in calibrated OM score, potentially through competing mechanisms. Cluster 2, in contrast, shows significant enrichment of a different set of taxa, including *Alloprevotella, Gemella*, and, *Prevotella*, while *Rothia, Actinomyces*, and *Neisseria* were relatively depleted. This phenotype is partially aligned with Ganly et al (2019), where *Prevotella, Alloprevotella, Fusobacterium*, and *Veillonella* were found to contribute to the enrichment of proinflammatory genes toll-like receptor (TLR) 1, TLR2, and TLR4. These particular microbial signatures characterizing Cluster 2 may possibly explain the steep increase in calibrated OM score. Cluster 3, displayed a significant upregulation of *Prevotella* and *Alloprevotella*, substantial evidence has shown that these two closely related genera contribute to oral inflammatory processes (Könönen et al, 2022; Pan et al,2023; Ganly et al, 2019) and are involved in systemic inflammatory conditions (Wolff et al, 2017). Meanwhile, Cluster 3 is also characterized by the significant depletion of *Stomatobaculum, Porphyromonas, Haemophilus*, and *Actinomyces*. Although the loss of oral *pathobiont Porphyromonas* may appear as beneficial, the concurring depletion of the other commensal taxa may contribute to calibrated OM score progression in cluster. Collectively, these cluster-specific microbial signatures highlight how distinct dysbiotic trajectories may drive either gradual or acute OM progression, with *Prevotella*-centered signatures emerging as a common feature of imbalances across the different clusters.

This study refines the findings of Zhang et al (2024), who previously examined longitudinal patterns of OM severity. We improved the statistical analyses of the longitudinal oral microbiome data in several key ways. First, by incorporating a broader set of self-reported symptom scores, we derived a more comprehensive and clinically informative OM score for patients with HNSCC. This new calibrated OM score was positively correlated with the OM score used by Zhang et al (2024) but was less noisy. Second, our patient clustering was based on a functional data approach and the selection of the number of clusters was based on the gap statistic (Tibshirani et al, 2001).

As a result, this approach improved interpretability compared to Zhang et al (2024), producing more distinctive microbial signatures. Finally, to identify statistically significant cluster-specific microbial signatures, we used PLSKO (Yang et al, 2025), which is a statistically principled procedure with appropriate false discovery control.

Given the differential microbial signatures identified in OM patient clusters, microbiome-based interventions present a promising avenue for managing OM severity and progression. Probiotic therapy could potentially help restore microbial balance and promote mucosal healing. Another potential approach is the targeted use of antimicrobial agents that could selectively suppress bacterial taxa overrepresented in severe OM cases. Further research into personalized microbiome modulation could provide innovative and personal therapeutic strategies for OM in HNSCC patients.

## Declarations

### Consent for publication

Not applicable

### Data availability

Available upon request.

### Materials availability

Not applicable

### Code availability

All analyses were conducted in R and available at https://github.com/SarithaKodikara/OMScoreAnalysis

### Author contribution

Conceived and designed the study: JM, SK. Performed the analysis: JM, SK. Wrote the paper: JM, SK, EMDSV, KAD, CCRG, KAL C.

## References

Aghabozorgi S, Seyed Shirkhorshidi A, Ying Wah T (2015) Time-series clustering – a decade review. Inf Syst 53:16–38

Balducci L, Extermann M (2000) Management of cancer in the older person: a practical approach. The oncologist 5(3):224–237

Barber RF, Candès EJ (2015) Controlling the false discovery rate via knockoffs. Annals of Statistics 43(5):2055–2085

Bell A, Kasi A (2025) Oral mucositis. StatPearls 12

Bik EM, Long CD, Armitage GC, et al (2010) Bacterial diversity in the oral cavity of 10 healthy individuals. ISME Journal 4(8):962–974

Carda-Diéguez M, Rosier B, Lloret S, et al (2022) The tongue biofilm metatran-scriptome identifies metabolic pathways associated with the presence or absence of halitosis. NPJ Biofilms and Microbiomes 8(1):100

Devaraja K, Aggarwal S (2025) Dysbiosis of oral microbiome: A key player in oral carcinogenesis? a critical review. Biomedicines 13(2):448

Ganly I, Yang L, Giese RA, et al (2019) Periodontal pathogens are a risk factor of oral cavity squamous cell carcinoma, independent of tobacco and alcohol and human papillomavirus. International Journal of Cancer 145(3):775–784

Ghaderi H, Foreman B, Nayebi A, et al (2024) Identifying tbi physiological states by clustering multivariate clinical time-series data. In: AMIA Annual Symposium Proceedings, p 379

Hampelska K, Jaworska MM, Babalska ZLt, et al (2020) The role of oral microbiota in intra-oral halitosis. Journal of Clinical Medicine 9(8):2484

Johnson DE, Burtness B, Leemans CR, et al (2020) Head and neck squamous cell carcinoma. Nat Rev Dis Primers 6(1):92

Kodikara S, Ellul S, Lê Cao KA (2022) Statistical challenges in longitudinal micro-biome data analysis. Brief Bioinform 23(4):bbac273

Könönen E, Fteita D, Gursoy UK, et al (2022) Prevotella species as oral residents and infectious agents with potential impact on systemic conditions. Journal of Oral Microbiology 14(1):2079814

Lê Cao KA, Welham ZM (2021) Multivariate Data Integration Using R, 1st edn. Chapman and Hall/CRC

Lê Cao KA, Boitard S, Besse P (2011) Sparse PLS discriminant analysis: biologically relevant feature selection and graphical displays for multiclass problems. BMC Bioinformatics 12(1):253

Li PJ, Li KX, Jin T, et al (2020) Predictive model and precaution for oral mucositis during chemo-radiotherapy in nasopharyngeal carcinoma patients. Frontiers in Oncology 10:596822

Lima KM, Alves CMC, Vidal FCB, et al (2023) Fusobacterium nucleatum and pre-votella in women with periodontitis and preterm birth. Medicina Oral, Patología Oral y Cirugía Bucal 28(5):e450

Lionel D, Christophe L, Marc A, et al (2006) Oral mucositis induced by anti-cancer treatments: physiopathology and treatments. Therapeutics and Clinical Risk Management 2(2):159–168

Marron JS, Dryden IL (2021) Object Oriented Data Analysis, 1st edn. Chapman & Hall/CRC, Philadelphia, PA

Pan J, Zhang X, Shi D, et al (2023) Correlation of oral microbiota with different immune responses to antiretroviral therapy in people living with HIV. Infectious Microbes & Diseases pp 10–1097

Peterson CB, Saha S, Do KA (2024) Analysis of microbiome data. Annu Rev Stat Appl 11(1):483–504

Pulito C, Cristaudo A, Porta CL, et al (2020) Oral mucositis: the hidden side of cancer therapy. J Exp Clin Cancer Res 39(1):210

Reyes-Gibby CC, Wang J, Zhang L, et al (2020) Oral microbiome and onset of oral mucositis in patients with squamous cell carcinoma of the head and neck. Cancer 126(23):5124–5136

Rosenthal DI, Mendoza TR, Chambers MS, et al (2007) Measuring head and neck cancer symptom burden: the development and validation of the md anderson symptom inventory, head and neck module. Head & Neck: Journal for the Sciences and Specialties of the Head and Neck 29(10):923–931

San Valentin EMD, Do KA, Yeung SCJ, et al (2023) Attempts to understand oral mucositis in head and neck cancer patients through omics studies: A narrative review. Int J Mol Sci 24(23):16995

Sigg CD, Buhmann JM (2008) Expectation-maximization for sparse and non-negative PCA. In: Proceedings of the 25th international conference on Machine learning - ICML ‘08. ACM Press, New York, New York, USA

Sonis S (1998) Mucositis as a biological process: a new hypothesis for the development of chemotherapy-induced stomatotoxicity. Oral oncology 34(1):39–43

Tibshirani R (1996) Regression shrinkage and selection via the lasso. J R Stat Soc Series B Stat Methodol 58(1):267–288

Tibshirani R, Walther G, Hastie T (2001) Estimating the number of clusters in a data set via the gap statistic. Journal of the Royal Statistical Society: Series B (Statistical Methodology) 63(2):411–423

Wold H (1966) Estimation of principal components and related models by iterative least squares. Multivariate Analysis pp 391–420

Wolff B, Boutin S, Lorenz H, et al (2017) Fri0698 prevotella and alloprevotella species characterize the oral microbiome of early rheumatoid arthritis. Annals of the Rheumatic Diseases 76:754

Yang G, Menkhorst E, Dimitriadis E, et al (2025) PLSKO: a robust knockoff generator to control false discovery rate in omics variable selection. Bioinformatics

Zhang L, San Valentin EMD, John TM, et al (2024) Influence of oral microbiome on longitudinal patterns of oral mucositis severity in patients with squamous cell carcinoma of the head and neck. Cancer 130(1):150–161

